# Tead1 reciprocally regulates adult β-cell proliferation and function to maintain glucose homeostasis

**DOI:** 10.1101/2020.03.05.979450

**Authors:** Jeongkyung Lee, Ruya Liu, Byung S. Kim, Yiqun Zhang, Feng Li, Rajaganapti Jagannathan, Ping Yang, Vinny Negi, Joseph Danvers, Eliana Melissa Perez-Garcia, Pradip K. Saha, Omaima Sabek, Chad J. Creighton, Cristian Coarfa, Mark O. Huising, Hung-Ping Shih, Rita Bottino, Ke Ma, Mousumi Moulik, Vijay K. Yechoor

**Author notes:** Correspondence should be addressed to V.K.Y. or J.L.

## Abstract

Proliferative quiescence in β-cells is required to maintain functional competence. While this presents a significant hurdle in regenerative therapy for diabetes, the molecular underpinnings of this reciprocal relationship remain unclear. Here, we demonstrate that TEAD1, the transcription effector of the mammalian-Hippo pathway, drives developmental stage-specific β-cell proliferative capacity in conjunction with its functional maturation. TEAD1 promotes adult β-cell mature identity by direct transcriptional control of a network of critical β-cell transcription factors, including, *Pdx1, Nkx6.1, and MafA,* while its regulation of *Cdkn2a* maintains proliferative quiescence. Consequently, mice with either constitutive or inducible genetic deletion of *TEAD1* in β-cells developed overt diabetes due to a severe loss of secretory function despite induction of proliferation. Furthermore, we show that TEAD1 has a similar regulatory role in human β-cells. Consistent with this function in β-cells, variants in *TEAD1* have been associated with c-HOMA-B in American Indians. We propose that TEAD1 is an essential intrinsic molecular switch coordinating adult β-cell proliferative quiescence with mature identity and its differential modulation may be necessary to overcome the challenge of inducing proliferation with functional competence in human beta cells.

---

Diabetes has assumed epidemic proportions with over 422 million people affected by it worldwide ^1^. A reduction of functional β-cell mass underlies all forms of diabetes. In type 1 diabetes, there is an autoimmune destruction of insulin-producing β-cells, while type 2 diabetes (T2D) is characterized by significant insulin resistance with varying degrees of reduction in β-cell mass and function ^2^. Current strategies to promote proliferative capacity in adult β-cells are often limited by the concurrent loss of mature function ^3-5^. Hence, a better understanding of the balance between proliferation and functional competence in mature β-cells is critical to enhancing functional β-cell mass to prevent and treat diabetes.

Tead family of transcription factors (Tead1-4) and their upstream modulators of the mammalian Hippo-Tead pathway are important regulators of cell proliferation to control organ size in embryonic development and tissue remodeling ^6-12^. Tead1, a canonical downstream transcriptional effector of the Hippo pathway is expressed ubiquitously, but its physiological function in mature differentiated cells is largely unknown. The Hippo signaling pathway consists of a core kinase cascade involving Mst1&2 kinases that phosphorylate Lats1&2, which in turn phosphorylate coactivators, Yap and Taz. Phosphorylated Yap/Taz are retained in the cytoplasm and subsequently targeted for degradation. In the absence of this kinase signaling Yap and Taz remain dephosphorylated, translocate to the nucleus and bind to Tead to regulate transcription ^11,13^. Interestingly, although various Hippo signaling components have been studied in pancreas biology, the role of Tead transcription factors in β-cells is unknown. Deletion of Mst1/2 in pancreatic progenitors resulted in Yap activation, ductal metaplasia and pancreatitis without overt β-cell defects in proliferation or function ^14,15^. During early pancreatic development Yap-Tead1 activate transcriptional events to control pancreatic progenitor ^16,17^, while Yap functions as an inhibitor of endocrine lineage specification, both in vivo ^18^ and during differentiation of iPS cells into β-cells ^19^. In contrast, though Yap is normally absent in adult β-cells ^14,15,20^, its overexpression promotes β-cell proliferation in human islets ex vivo ^21,22^, while CTGF/CCN2, a transcriptional target of Hippo-Tead1 pathway, also regulates β-cell proliferation ^23,24^. All this strongly suggest that the Hippo-Tead1 pathway regulates β-cell proliferation. However, the role of Tead1 in adult β-cell proliferation and function has remained unexplored.

In this study, employing cell-type and developmental stage-specific genetic tools to systematically probe Tead1 function in β-cells, we uncover its surprising role in driving mature β-cell identity and glucose stimulated insulin secretion (GSIS), while maintaining proliferative quiescence by direct transcriptional activation of a network of critical transcription factors and genes that confer functional competence and limit proliferation. Our study thus identifies Tead1, the Hippo effector, as a non-redundant reciprocal regulator of the proliferative drive and functional competence in β-cells. Differential modulation of these activities may allow an increase in functional β-cell mass without a loss of critical glucose homeostatic function as a therapeutic approach to diabetes.

## Results

### Tead1 is expressed in embryonic and adult beta cells

Tead family of transcription factors regulate proliferative pathways and determine organ size in many tissues during development ^11,25^. However, their role in β-cell development and function are unknown. Examination of islets at different developmental stages revealed that Tead1 was preferentially expressed in adult mouse islets, and highly enriched as compared to other members of the Tead family that are present at 10-fold lower level (Tead2) or not detected (Tead 3 &4) (**Figure. 1A**). Of the two canonical Tead1 co-activators Yap and Taz, only Taz was robustly expressed (**Figure. 1A**), as were many upstream signaling components of Hippo pathway (Mst1&2, Lats1&2 and others), suggesting the Hippo-Tead pathway is functional in islets. Expression dynamics of Tead1 by immunostaining during islet development revealed it was most abundant at e15.5 in all pancreatic tissues, but its high expression was restricted to islets around weaning and persisted at a high level in adult β-cells (**Figure. 1B**). While Taz was robustly expressed throughout, from e15.5 into adult stage, Yap was absent in the islets at all stages examined (**Figure. 1B**), a finding that is consistent with previous studies ^14^. Human islets also expressed robust levels of TEAD1 and Taz but not YAP **(Figure 1C)**. In contrast to mouse islets, human islets express higher levels of Tead4. Based on the robust, yet restricted expression of Tead1 in mature β-cells, we selectively deleted Tead1 in a developmental stage-specific manner by crossing mice with Tead1 floxed (Tead1^F/F^) alleles carrying a floxed exon 3 that encodes the DNA binding TEA domain (**Figure S1A-C**) ^26^ with three distinct Cre lines, namely Rip-Cre, Ins1-Cre (Thorens), and Mip-CreERT.

**Figure 1:**
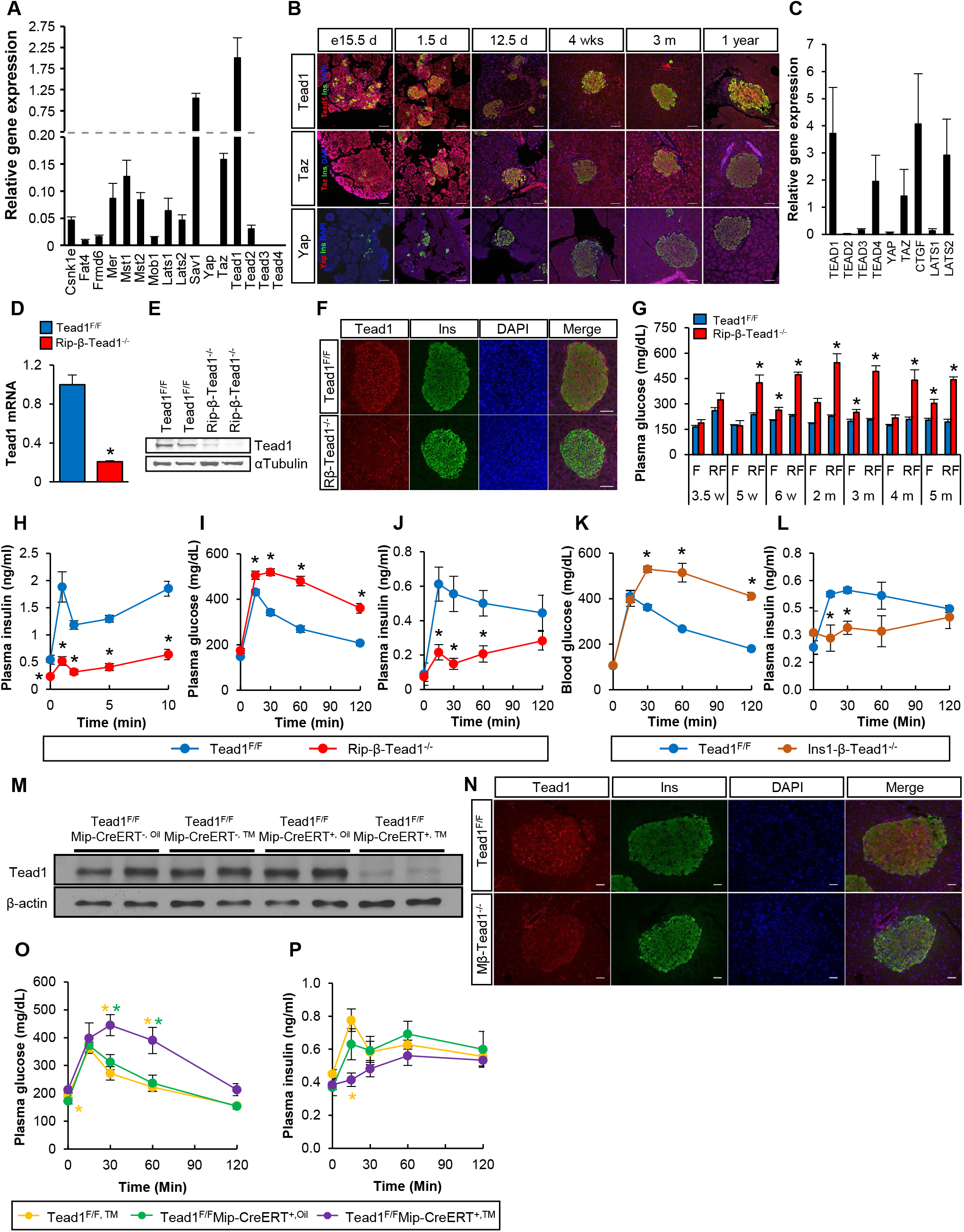
β-cell Tead1 deletion leads to diabetes. (A) Relative gene expression assessed by RT-qPCR of total RNA from isolated islets of 4 month old wild type mice, normalized to housekeeping gene, *Tbp*. n=4 (B) Representative immunofluorescence staining of Tead1, Taz and Yap in mouse islets at different ages. Scale bar - 50µm. (C) Relative gene expression assessed by RT-qPCR of total RNA from isolated human islets, normalized to housekeeping gene, *TBP.* n=4 (D) RT-qPCR (normalized to control Tead1^F/F^) n=4-6; and (E) western blotting for Tead1 in isolated islets from Tead1^F/F^ and Rip-β-Tead1^-/-^mice. (F) Representative images of immunofluorescence staining of Tead1^F/F^ and Rip-β-Tead1^-/-^ pancreas. Scale bar - 50µm. (G) Plasma glucose level with overnight fasting and after 1 hour refeeding in Tead1^F/F^ and Rip-β-Tead1^-/-^mice. n=4-6. w-weeks; M-months; F-fasting; RF- refeeding. (H) Acute insulin secretion after glucose stimulation in 10-week-old mice. n=5. (I) Plasma glucose and (J) insulin during glucose tolerance testing (GTT) in overnight fasted 8 weeks old male mice. n=5-6. (K) Plasma glucose and (L) insulin during GTT in overnight fasted 9 weeks old male mice. n=3-4. (M) Western blotting of isolated islets from Tead1^F/F^ and Tead1^F/F^Mip-CreERT^+^ mice treated with oil or Tamoxifen (TM). (N) Nuclear Tead1 staining is absent from most insulin expressing β-cell in TM-treated Mip-β-Tead1^-/-^ islets. Scale bar – 50µm. (O) Plasma glucose level and (P) insulin levels during the GTT in overnight fasted 5 month old Tead1^F/F^ and Tead1^F/F^Mip-CreERT^+^ (Mip-β-Tead1^-/-^) mice treated with oil or TM. n=4. TM, Tamoxifen. All values are mean ± SEM, * - p≤ 0.05.

### Tead1 deletion in β-cells leads to diabetes

Early constitutive deletion at e15.5 (Rip-β-Tead1^-/-^), using Rip-Cre deletor mice, led to a loss of β-cell Tead1 expression, as indicated by >80% reduction in transcript (**Figure 1D**) and near absence Tead1 protein (**Figure 1E**) in whole islets. Loss of Tead1 immunostaining was specific to β-cells (**Figure 1F**), but not other hormone-producing islet cells (data not shown). These Rip-β-Tead1^-/-^ mice with embryonic deletion of Tead1 developed progressive hyperglycemia (fasting and fed) within 2 weeks of weaning at 5 weeks of age. At 2 months of age, they became diabetic with fed plasma glucose >500mg/dl (**Figure 1G**). Indicative of a severe defect in β-cell secretory function, the Rip-β-Tead1^-/-^ mice lacked the first phase insulin secretion (**Figure 1H)**, a feature that is lost very early in type 2 diabetes ^27,28^, and displayed marked glucose intolerance during an intraperitoneal glucose tolerance testing (GTT) due to a blunted GSIS (**Figure 1I-J**). These results were not secondary to changes in body weight (**Figure S1D**) or insulin sensitivity (**Figure S1E**). Overall energy balance was similar to controls as indicated by food intake and whole body energy expenditure, VC02, V02, and RER (**Figure S1F-J**). To exclude potential confounding effect of hypothalamic expression or potential secretory dysfunction due to the Rip-Cre transgene ^29,30^, we applied another Ins1-Cre deletor line ^31^ for specific genetic targeting of Tead1 in β-cells by generating the Ins1-β-Tead1^-/-^mice. These Ins1-β-Tead1^-/-^ phenocopied the Rip-β-Tead1^-/-^ indicated by marked glucose intolerance with a significant blunting of GSIS in vivo during GTT **(Figure 1K-L).**

While the findings in Ins1-β-Tead1^-/-^ and Rip-β-Tead1^-/-^mice demonstrated the requirement of Tead1 function in β-cells in maintaining insulin secretion and glucose homeostasis in vivo, these models that involve embryonic deletion starting at e15.5 could also engender chronic compensatory changes. Hence, we tested whether Tead1 modulates adult β-cell function using Mip-CreERT mice ^29^ to achieve tamoxifen (TM)-inducible deletion (Mip-β-Tead1^-/-^ mice). Tamoxifen administration (100mg/kg every other day for 5 doses by gavage) in 5-week-old Mip-β-Tead1^F/F^ mice largely abolished-Tead1 protein expression in whole islet lysates when examined after 6 weeks, (**Figure 1M**), along with an absence of Tead1 nuclear immunostaining in ∼85% of β-cells (**Figure 1N**). Strikingly, this restricted inactivation of Tead1 only in mature β-cells led to a marked glucose intolerance due to a blunting of in vivo GSIS with a loss of early phase insulin secretion (**Figure 1O-P**), demonstrating severe β-cell secretory deficits within 2 months post-TM treatment. Taken together, the three loss-of-function models of Tead1, during β-cell development (Rip-β-Tead1^-/-^ and Ins1-β-Tead1^-/-^) or after maturation (Mip-β-Tead1^-/-^) demonstrate the essential role of Tead1 in maintaining β-cell secretory capacity and glucose homeostasis.

### Tead1 transcriptional control of a regulatory network is critical for maintaining functional mature β-cell identity

Using isolated Rip-β-Tead1^-/-^ islets, we next determined the specific functional deficits induced by a loss-of-Tead1 function. Despite normal basal insulin secretion these islets display significantly impaired GSIS with a markedly lower insulin secretion index (**Figure 2A**), as compared to either Floxed or Rip-Cre controls. Furthermore, in Ins-1 (832/13) cells, Tead1 knockdown yielded a similar blunting of GSIS response (**Figure S2A-B**), suggesting the cell-autonomous role of Tead1 for GSIS in β-cells. Consistent with the GSIS deficits, there were ∼50-80% reductions of insulin content in Rip-β-Tead1^-/-^ mice, whether measured in the whole pancreas (**Figure 2B**), normalized to islet size **(Figure 2C**) or by cell number (reflected by DNA content) as compared to controls (**Figure 2D**).

**Figure 2:**
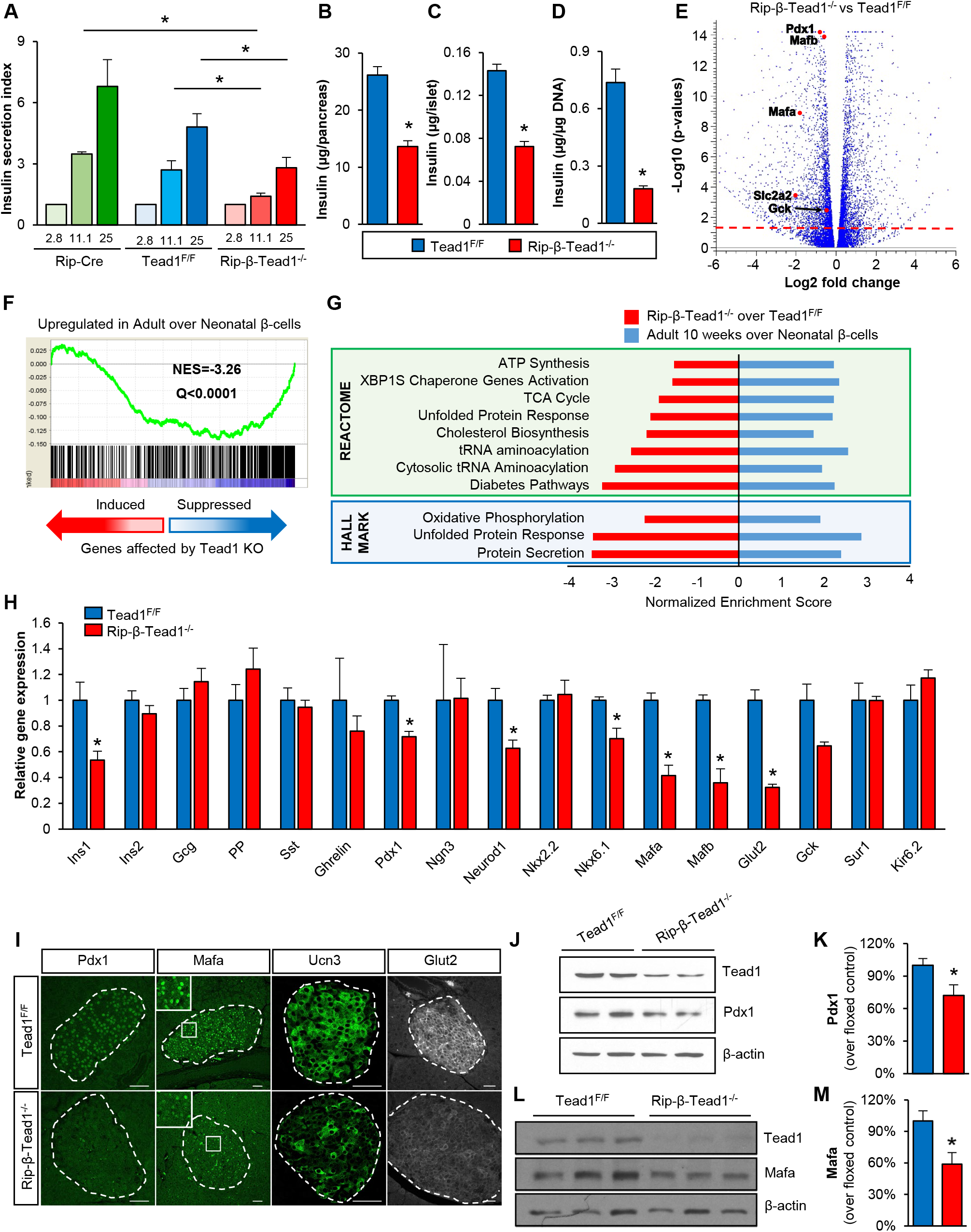
Tead1 is required for mature β-cell function. (A) GSIS from isolated islets (ten islets each from 5-6 individual mice of each genotype) on exposure to increasing glucose concentrations. Insulin secretion index is expressed as a fold change over basal secretion in 2.8mM glucose. Insulin content in (B) total pancreas, n=4; in isolated islets expressed as insulin content (C) per equally sized islet or (D) per μg DNA, n=5. (E) Volcano plot from RNA-seq data from differentially expressed genes from isolated islets of 1 year old Rip-β-Tead1^-/-^ compared to Tead1^F/F^. Some of the genes critical in mature β-cell function are highlighted. (F) GSEA analysis of the differentially expressed downregulated genes in isolated islets of Rip-β-Tead1^-/-^ compared to Tead1^F/F^ as compared to the published gene set (GSE47174) that was upregulated in adult β-cells when compared to neonatal β-cells. Details in the accompanying text. NES – normalized enrichment score. FDR Q value is also shown. (G) Enriched pathways in transcriptome analysis of Rip-β-Tead1^-/-^ compared to Tead1^F/F^ isolated islets. x-axis shows the Normalized enrichment score. Note that in this representation the red bars indicate the enrichment of Rip-β-Tead1^-/-^ over control Tead1^F/F^ indicating these pathways are suppressed in islets from Rip-β-Tead1^-/-^ mice, while the blue bars indicate that the same pathways are upregulated in adult over neonatal β-cells. (H) Expression, by RT-qPCR, of genes critical to mature β-cell function in Tead1^F/F^ and Rip-β-Tead1^-/-^ islets, normalized to housekeeping gene, *TopI.* n=4. (I) Representative immunofluorescent images of Pdx1, MafA, Ucn3 and Glut2 staining in Tead1^F/F^ and Rip-β-Tead1^-/-^ islets. Scale bar - 50 μm. (J-M) Western blot and quantification of (J and K) Pdx1 protein and (L and M) MafA from isolated islets from Rip-β-Tead1^-/-^ and Tead1^F/F^ mice. All values are mean ± SEM, * - p≤ 0.05.

To determine the regulatory pathways mediating Tead1 function in islets, we performed global transcriptome analysis in Rip-β-Tead1^-/-^ islets that revealed 259 (down) and 296 (up) differentially regulated genes as compared to age-matched floxed controls (**Figure S2C and Supplementary table 3**). In agreement with the profound defects in GSIS, among the differentially regulated genes by RNA-seq analysis distinct components of the insulin secretory pathway including the glucose transported Glut2 (Slc2a2) and glucokinase (Gck) were down-regulated in Tead1-deficient islets. In addition, key transcription factors known to specify and maintain adult β-cell function were suppressed, such as Pdx1, MafA. (**Figure 2E**). Further analysis using an unbiased Gene Set Enrichment Analysis (GSEA) revealed a global down-regulation in adult β-Tead1^-/-^ islets of genes significantly enriched in mature adult as compared to immature neonatal islets (GSE47174) ^32^ (**Figure 2F)**. Interestingly, many disallowed genes in normal adult β-cells that comprise the neonatal islet transcriptome signature ^33-36^ and are highly expressed in immature neonatal islets were not altered in Tead-1 deficient islets (**Figure S2D**). These findings suggest a loss of mature β-cell characteristics, rather than de-differentiation ^5^ of Tead1-null β-cells. Further support of this notion was found in the transcriptomic signatures of key processes that are critical for acquisition and maintenance of mature secretory function in β-cells, including OXPHOS, substrate metabolism, protein secretion and export, vesicular transport and UPR were consistently suppressed with loss of Tead1 (GSEA, q<0.25) **(Figure 2G**). We further validated that in Rip-β-Tead1^-/-^ islets, transcript (**Figure 2H)** of Ins1, Glut2, Pdx1, NeuroD, MafA and Nkx6.1 were all significantly decreased along with Ucn3, a hallmark of mature β-cells ^37,38^. The protein levels of many of these genes were also significantly reduced by immunostaining (**Figure 2I**) or western blotting **(Figure 2J & 2L)** with respective quantifications (**Figure 2K & 2M**). These data collectively demonstrated Tead1 transcriptional control of a regulatory network critical for maintaining functional mature β-cell identity.

We interrogated whether Tead1 exerts direct transcriptional regulation of this gene network driving β-cell function and identity. Analysis of Tead1 occupancy by ChIP-seq in adult mouse islets on regulatory regions of transcription factors that are critical for β-cell function, including Pdx1, MafA, Nkx6.1, revealed its enrichment on these genes that were downregulated by loss of Tead1 (**Figure 3A**). ATAC-seq further confirmed that the identified promoter regions with Tead1 occupancy in islets occurred in open chromatin regions that are transcriptionally active (**Figure 3A)**. ChIP-qPCR using either endogenous Tead1 (**Figure 3B**) or tagged Myc-Tead1 (**Figure S3A**) in mouse insulinoma cell line further validated Tead1 binding to cis-promoter elements (containing Tead consensus binding sequences of GGAATG) of Pdx1, MafA, Nkx6.1, Ucn3.

**Figure 3:**
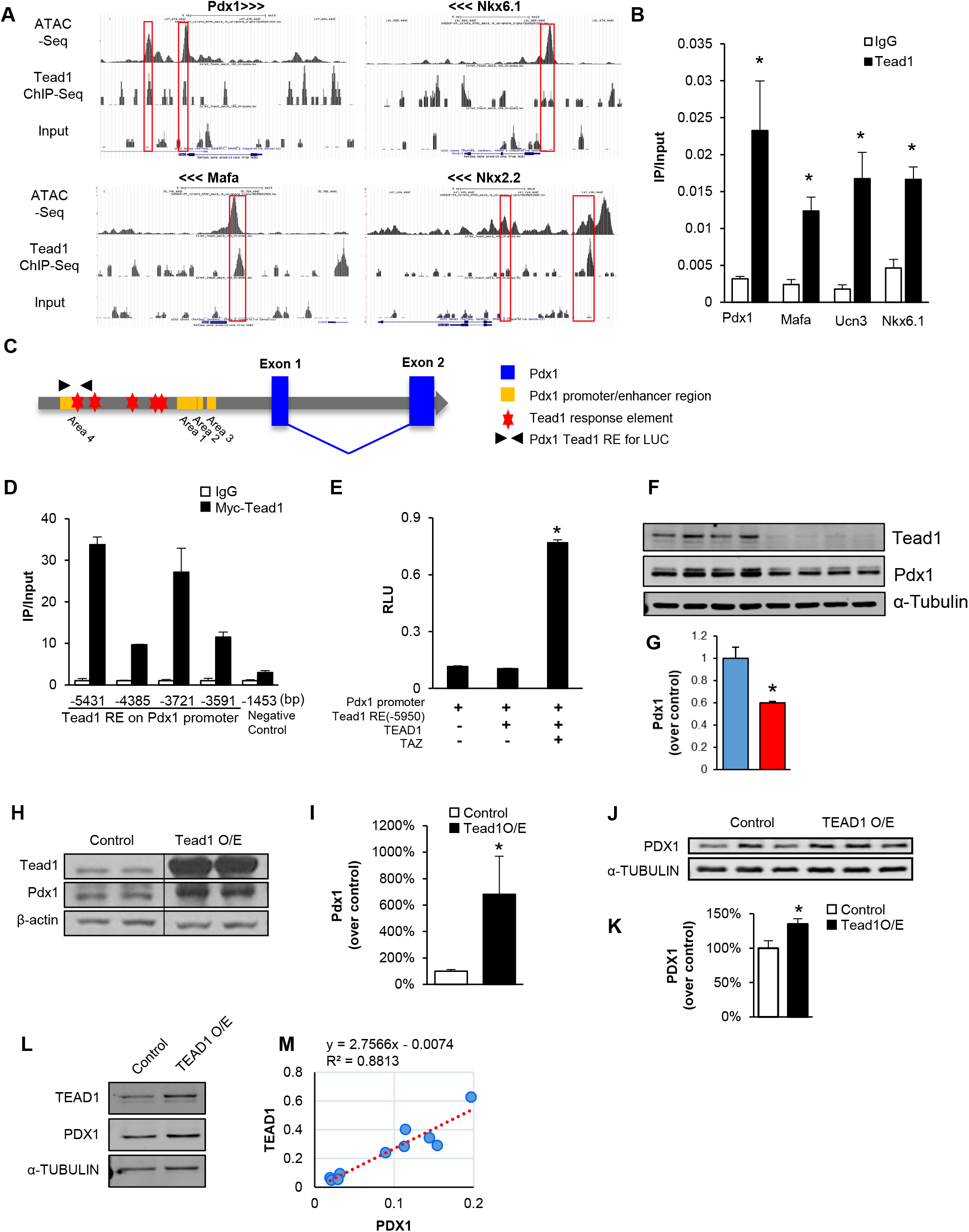
Tead1 is direct transcriptional regulator of genes for mature β-cell function. (A) Tead1 occupancy in the active promoters of displayed genes that have open chromatin in mouse islets by ATAC-seq and ChIP-seq. Red boxes indicate overlapping regions of Tead1 occupancy (Tead1 Chip-seq) and open chromatin (ATAC-seq). Input control is also shown. (B) ChIP in Ins-2 cells with Tead1 or control IgG antibody and qPCR with primers flanking putative Tead1 response elements. The y-axis represents the ratio of pulldown DNA to input DNA. n=3. (C) Tead1 response elements (Tead1 RE) in promoter region of mouse *Pdx1* is shown schematically and (D) ChIP in Myc-Tead1 over-expressed Ins-2 cells with Myc or control IgG antibody and qPCR with putative Tead1 response element on mouse Pdx1 promoter. (E) Promoter luciferase assay with native Pdx1 promoter-Tead1 response element (RE:Luc) reporter in Ins-2 cells. y-axis represents the relative luminescence units (RLU). n=3. (F-G) Western blotting for Pdx1 protein in Tead1 knock down Ins2 cell line with quantification relative to α-tubulin as housekeeping control (G). (H-I) Western blotting and quantification for PDX1 protein in TEAD1 overexpressing (Tead1 O/E) in mouse Ins2 cell line, in (J and K) human EndoC-β-h2 cell line and in (L) human islets. Control cells are transfected with empty vector. (M) Correlation analysis of PDX1 protein and TEAD1 in human islets (n=8). All values are mean ± the standard errors of the means, * - p≤ 0.05.

Since Pdx1 is a master regulator of mature β-cell function, we undertook a detailed analysis of Tead1 regulation of Pdx1 transcription. Interestingly, we found multiple Tead1 responsive elements in the Pdx1 promoter region as represented in **Figure 3C**, including the Tead1 consensus sequence present in Area IV region known to be critical for β-cell specific Pdx1 expression and postnatal mature β-cell function ^39^. In ChIP pull down with Myc-Tead1 in Ins2 cells, this site exhibited the highest enrichment (**Figure 3D**). In a promoter luciferase assay, expression of Tead1 together with its co-activator, Taz, activated transcription of the native Pdx1-promoter containing Tead1 response elements, demonstrating Tead1 functional regulation of Pdx1 transcription (**Figure 3E**). Furthermore, Tead1 knockdown decreased Pdx1 protein level in Ins2 cells (**Figure 3F-G)**; whereas its overexpression by adenoviral vector in Ins2 cells increased Pdx1 expression (**Figure 3H-I**). Overexpression of TEAD1 in additional human β-cell models including human β-cell line (EndoC-βh2) **(Figure 3J-K**) or human cadaveric islets (**Figure 3L**) augmented PDX1 expression. It is noteworthy that TEAD1 and PDX1 protein levels detected by western blotting in individual human islet samples exhibited significant linear correlation (**Figure 3M)**. Thus, TEAD1 transcriptional control of PDX1 is conserved in human β-cells. Taken together, these data provide direct support of Tead1 as a direct transcriptional regulator that orchestrates a transcription factor network regulatory network critical to acquire and maintain mature β-cell function.

### Tead1 mediates proliferative quiescence in adult β-cells

Tead1 drives cell proliferation in various tissue development processes ^11-13^. We postulated that Tead1 deficiency during islet development may limit β-cell proliferation to reduce β-cell mass. Surprisingly, Rip-β-Tead1^-/-^ islets from 12 week old mice contained increased number of β-cells (**Figure 4A**), which was also reflected by increased DNA content in similar-sized islets (**Figure S4A**). However, these β-cells were significantly smaller (**Figure 4B-C)**, which offset the increase in numbers, so β-cell mass in Rip-β-Tead1^-/-^ mice remained normal compared to that of the floxed controls (**Figure 4D**). Similar findings were observed when these were repeated in 5-6 month old Ins1-β-Tead1^-/-^ islets (**Figure S4B-D**). Tead1 deletion only in adult β-cells in the inducible line of 9 month old adult Mip-β-Tead1^-/-^ mice yielded similar findings with a significant decrease in single β-cell area (**Figure S4E**), but resulted in no change in total β-cell area (**Figure S4F**) or islet size (**Figure S4G**). Further analysis indicated that increased β-cell number in Rip-β-Tead1^-/-^ islets was not due to apoptosis (**Figure S4H-I**), but rather because of higher proliferation rate, as reflected by a significantly elevated Ki67 positive β-cells (2% vs. 0.62% in 12 week old Rip-β-Tead1^-/-^ and Tead1^F/F^ controls, respectively; p<0.01) (**Figure 4E-F**).

**Figure 4:**
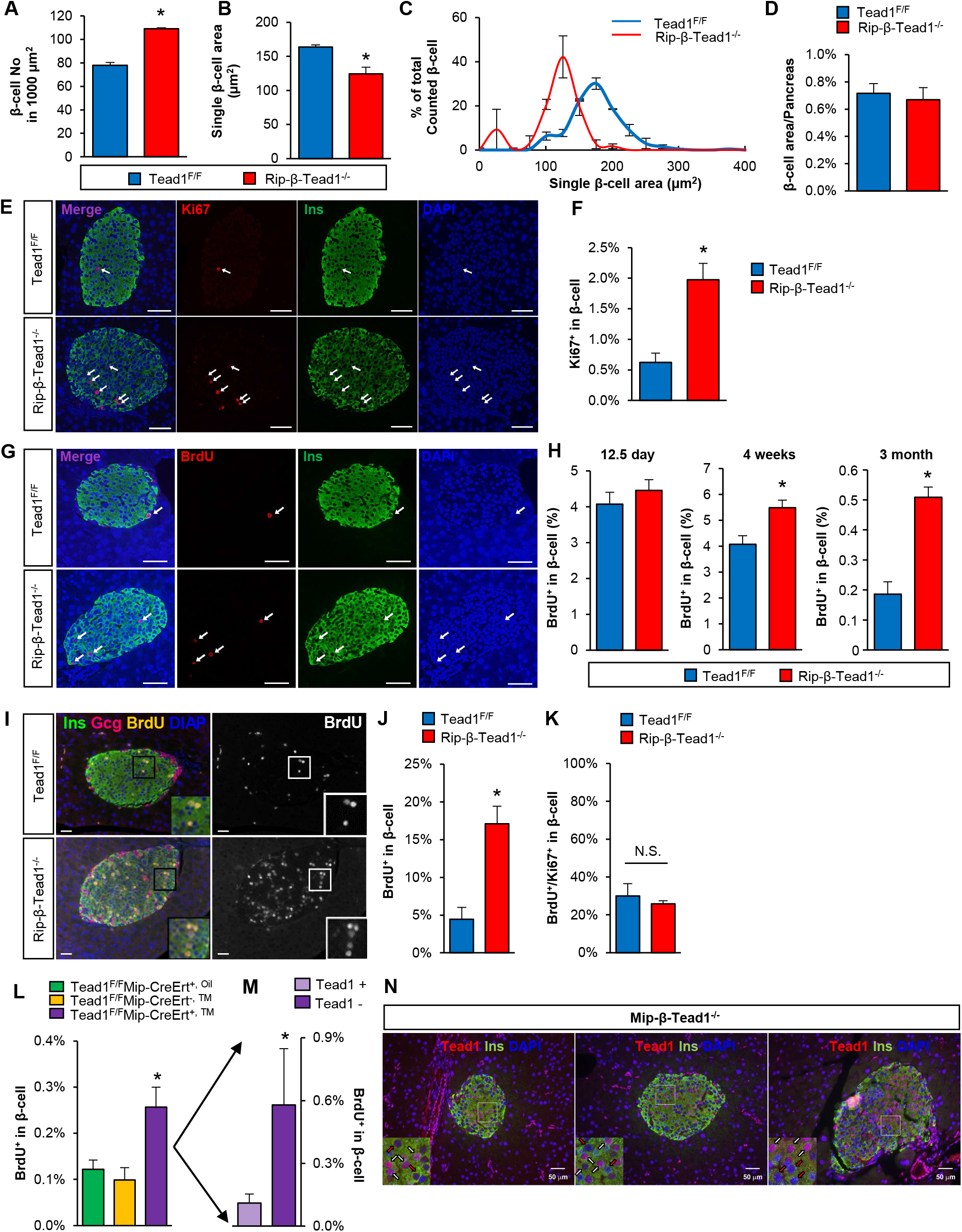
Tead1 deletion enhances β-cell proliferation. (A) β-cell number in 1000 µm^2^ in Rip-β-Tead1^-/-^ mice and Tead1^F/F^ mouse islet. N=3. (B) Single β-cell area in Rip-β-Tead1^-/-^ mice and Tead1^F/F^ mouse pancreas at 3 months age. n=3. (C) β-cell size distribution in Tead1^F/F^ mice (blue) and Rip-β-Tead1^-/-^ mice (red) at 3 month age. All values are means ± SEM. n=3. (D) Total pancreatic β-cell area from Rip-β-Tead1^-/-^ mice and Tead1^F/F^ mice at 3 months age. n=3. (E-H) Quantification of β-cell proliferation in Tead1^F/F^ and Rip-β-Tead1^-/-^ mice at 3 months. n=3. BrdU was injected (G and H) 2 hrs prior to sacrifice, or (I and J) administered continuously by water for 6 days prior to sacrifice. Representative microscopy images of Rip-β-Tead1^-/-^ mice and Tead1^F/F^ mouse pancreas at 3 months age and quantification of (E and F) Ki67 expressing and (G and I) BrdU positive β-cells. n=4. Scale bar – 50µm. (K) The fraction of BrdU-positive β-cells to Ki67-positive β-cells in the Rip-β-Tead1^-/-^ and controls. n=3-4 (L) Quantification of Brdu-positive β-cells in in 11 month old TM-treated Mip-β-Tead1^-/-^ mice and controls. n=4. (M) Brdu-positive cells in Tead-negative and Tead1-positive β-cell subpopulations in 11 month old TM-treated Mip-β-Tead1^-/-^ mice. n=4. All values are mean ±the standard errors of the means, * - p ≤ 0.05. N.S.-not significant. (N) Mosaic deletion of Tead1 in TM-treated Mip-β-Tead1^-/-^ β-cells leads to intra-islet mosaicism of Tead1 expression in β-cells. Representative confocal microscopy images of immunofluorescent staining of pancreas sections from Mip-β-Tead1^-/-^ mice indicating Tead1-neg (red arrow) and Tead1-pos (white arrow) β-cells in 3 different mice.

To test if Tead1 regulates β-cell proliferation at distinct developmental stages, we analyzed in vivo BrdU incorporation into β-cells in neonatal (postnatal day 12), post-weaning (4 weeks) and adult (12 weeks) mice. Notably, while there was no significant difference in proliferation at postnatal day 12 between Rip-β-Tead1^-/-^ and Tead1^F/F^ littermate controls, higher rate of BrdU incorporation was observed at 4 (5.31% vs. 4.08%; p<0.03) and 12 weeks of age (0.5% vs. 0.2%; p<0.05) respectively (**Figure 4G-H),** with similar increase (0.03% vs. 0.1%; p<0.05)seen in 5-6 month old Ins1-β-Tead1^-/-^ islets compared to floxed controls (**Figure S4D**). This data indicate that Tead1 did not contribute physiologic neonatal β-cell proliferative expansion when the proliferative drive is high, but rather was required for attaining proliferative quiescence as β-cells mature in adult mice.

β-cells have a short quiescence phase of ∼7 days after mitosis before they can re-enter cell cycle ^40^. Hence, we assessed if Tead1 could regulate total pool of actively proliferating β-cells by labelling all cycling β-cells over a 6 day period with BrdU via drinking water and found that active proliferation was ∼4 fold higher in Tead1-deficient β-cells than that of controls (17.1% vs. 4.46%) (**Figure 4I-J**). However, the fraction of proliferating β-cells in S-phase (BrdU-positive) to actively cycling Ki67-positive cells did not differ between the Rip-β-Tead1^-/-^ and controls (**Figure 4K**) indicating that loss of Tead1 did not affect cell cycle kinetics, once the cells entered cell cycle. Together, these data supports the notion that higher proliferation in Tead1-deficient β-cells likely occurred as a result of more cells entering cell cycle (G1) from the G0 quiescent state. We then asked if Tead1 affects age-associated proliferative quiescence when most β-cells become resistant to proliferative signals ^41^. Tead1 deletion in β-cells in 9 months old TM-treated Mip-β-Tead1^-/-^ mice showed a 2-fold increase in proliferation (**Figure 4L)**, similar as young adult mice. Thus, Tead1 was required to maintain quiescence in adult β-cells, while its inactivation releases cells from age-associated quiescence into active cell cycle.

The augmented β-cell proliferation in 4 week old Rip-β-Tead1^-/-^ mice (**Figure 4G**) preceded the onset of hyperglycemia at 5 weeks of age (**Figure 1F**), suggesting that the observed increase in proliferation is not secondary to hyperglycemia-induced compensation. However, to conclusively exclude non-cell autonomous effects, such as hyperglycemia, as a potential cause of proliferation, we leveraged the mosaic β-cell Tead1 deletion often seen with Mip-CreERT ^29,42^. Using this model, we pulse labelled cycling β-cells for 2 hours using BrdU and analyzed islets in which ∼50% of β-cells had deletion of Tead1. We found that within the same pool of islets in Mip-β-Tead1^-/-^ mice with Tead1-positive and Tead1-negative β-cells, the proliferative rate was ∼ 5 fold higher in Tead1-negative than that of the Tead1-positive β-cells (0.58% vs. 0.11%; p<0.05) (**Figure 4M-N**), indicative of cell-autonomous effect of Tead1 on proliferation.

### Tead1 is a direct transcriptional activator of p16 in adult β-cells

Corroborating the unexpected increased proliferation of Tead1-deficient β-cells, global transcriptome analysis of Tead1-deficient islets revealed significant upregulation of pathways involved in cell cycle control, including many G1-S checkpoint regulators (**Figure 5A**). Expression of Cyclin A2, critical for G1-S phase transition in β-cells ^43^ was significantly elevated in Rip-β-Tead1^-/-^ islets (**Figure 5B**). Though absence of Tead1 reduced Cyclin D1 transcript, its protein level was not altered (data not shown). In contrast, there were marked reductions of p16^INK4a^ and p19^ARF^, key cell cycle inhibitors critical for limiting β-cell proliferation in adult mice ^44-46^ (**Figure 5B**). p16^INK4a^ protein was nearly absent in Tead1-deficient islets (**Figure 5C-D**) as well as in Tead1-depleted Ins2 cells (**Figure 5E-F, and Figure 4F**), while Tead1 overexpression robustly increased p16^INK4a^ protein levels (**Figure 5G**). Examination of the *Cdkn2a* locus that encodes both p16^INK4a^ and p16^ARF^ demonstrated Tead1 occupancy in multiple Tead1 consensus motifs (**Figure 5H**). ChIP-qPCR in Ins2 cells demonstrated Tead1 binding to these motifs within the promoter of p16^INK4a^ in the *Cdkn2a* locus (**Figure 5I**). This was consistent with Chip-seq and ATAC-seq data from adult islets demonstrating Tead1 occupancy within Tead1 consensus motif containing regions of open chromatin in *Cdnk2a* locus (**Figure 5J**). Furthermore, Tead1 binding to these sites are functional as it can activate the native *Cdkn2a* (p16INK4a) promoter when co-expressed with Taz (**Figure 5K**).

**Figure 5:**
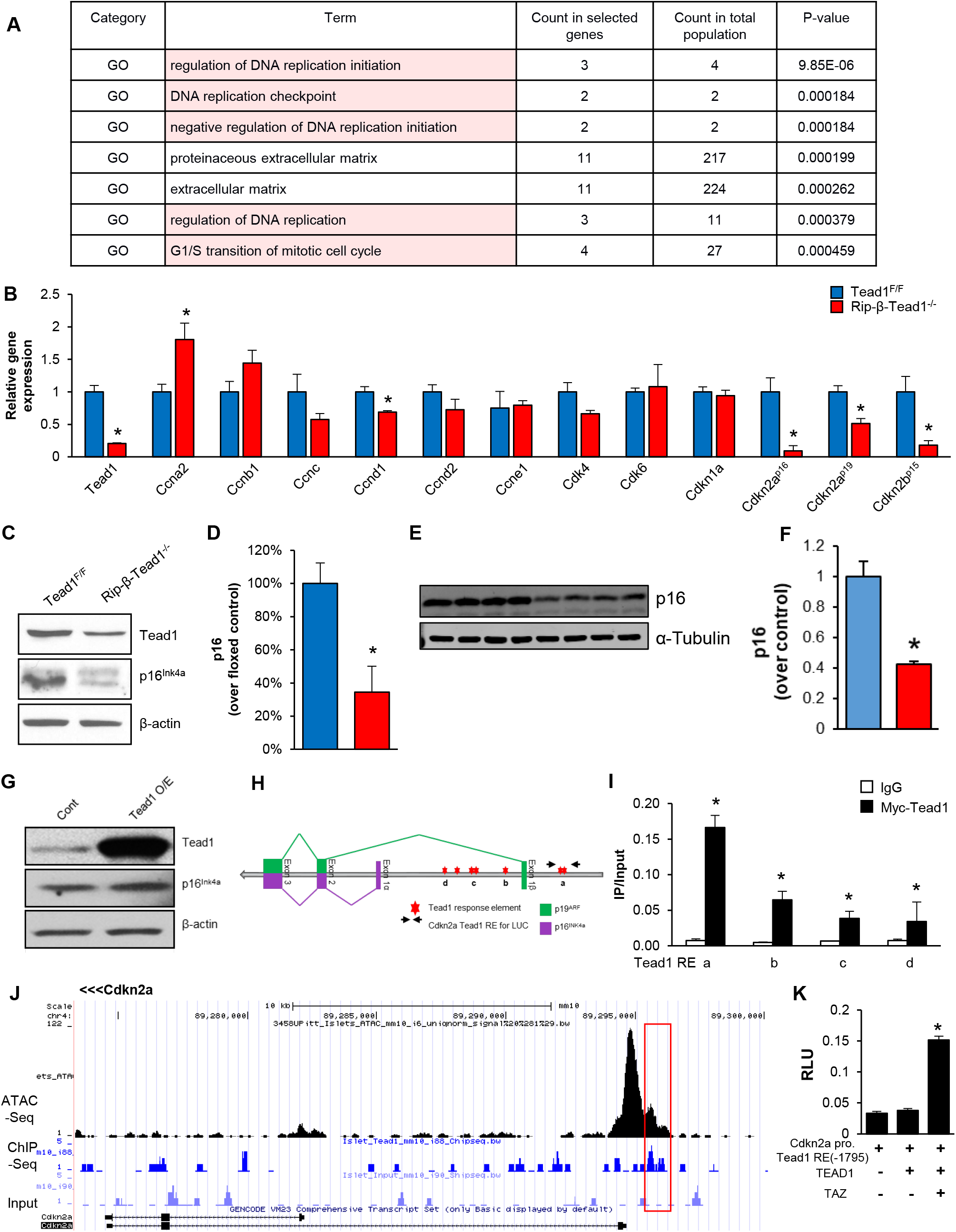
Tead1 is direct transcriptional regulator of genes for β-cell proliferation. (A) Transcriptome analysis of isolated Rip-β-Tead1^-/-^ and control Tead1^F/F^ islets from 12 weeks old male mice with significantly enriched pathways by Gene Ontology pathway analysis. All proliferation related pathways are highlighted in red. Details in the accompanying text. NES – normalized enrichment score. FDR Q value is also shown. (B) Relative expression of cell cycle related genes in Rip-β-Tead1^-/-^ and Tead1^F/F^ islets, normalized to housekeeping gene, *TopI*. n=4. (C) Western blot of p16^INK4a^ protein and (D) quantification in isolated islets from Rip-β-Tead1^-/-^ and Tead1^F/F^ mice. n=4-5. (E and F) Western blotting for p16^INK4a^ in (E) shRNA Tead1 knockdown and scrambled Ins-2 cells and quantification compared to α-tubulin as housekeeping control (F). This is from the same lysate as shown in Figure 3F. (G) Tead1 overexpression and control in Ins-2 cells. (H) Tead1 response elements (Tead1 RE) in promoter region of mouse *Cdkn2a*. The *Cdkn2a* locus encoding p16^INK4a^ and p19^ARF^ is shown schematically. (I) ChIP of Myc-Tead1 over-expressing Ins-2 cells with Myc or control IgG antibody and qPCR with primers flanking putative Tead1 response elements. The y-axis represents the ratio of pulldown DNA to input DNA. n=3. (J) Tead1 occupancy in the active promoter/intronic region of *Cdkn2a* that have open chromatin in mouse islets by ATAC-seq and ChIP-seq. Red box indicate overlapping regions of Tead1 occupancy (Tead1 Chip-seq) and open chromatin (ATAC-seq). Input control is also shown. (K) Promoter luciferase assay with native Cdkn2a promoter-Tead1 response element (RE:Luc) reporter in Ins-2 cells. y-axis represents the relative luminescence units (RLU). n=3. All values are mean ±the standard errors of the means, * - p≤ 0.05.

p16 represents a key cell cycle entry barrier to transition from a quiescent state, and Ezh2 has an established role as its critical repressor in β-cells ^41,45,46^. To explore whether Tead1 transcriptional regulation of p16^INK4a^ determines the balance between proliferation and quiescence, and if β-cell proliferation and its mature function were inter-dependent we tested if re-expressing p16 in Tead1-null β-cells in vivo through genetic ablation of its repressor Ezh2 could restore β-cell quiescence and insulin-secretory function. We inactivated Ezh2 in β-cells of Rip-β-Tead1^-/-^ mice (DKO) by crossing Ezh2 flox mice with Rip-β-Tead1^-/-^ mice. As expected of Ezh2 regulation of p16, the loss of Ezh2 in Tead1-null β-cells restored the level of p16 to a comparable level in wild type controls (**Figure 6A-B**). Notably, 6 month old Rip-β-Tead1^-/-^ β-cells without Ezh2 showed much lower proliferation rates than that seen in Rip-β-Tead1^-/-^ mice (Figure 4G-H) and at a level comparable to the control flox mice, (**Figure 6C-D**). Therefore, restoring p16 through the modulation of Ezh2 was sufficient to achieve/rescue proliferative quiescence in Tead1-null β-cells. Interestingly, the restoration of proliferative rates to normal was not sufficient in restoring glucose tolerance in these DKO mice, as these mice exhibited persistence glucose intolerance as compared to the floxed controls though there was some improvement as compared to littermate Rip-β-Tead1^-/-^ mice (**Figure 6E**). Thus, Tead1 regulation of proliferative quiescence was mediated by its direct transcriptional regulation of p16^INK4a^ expression.

**Figure 6:**
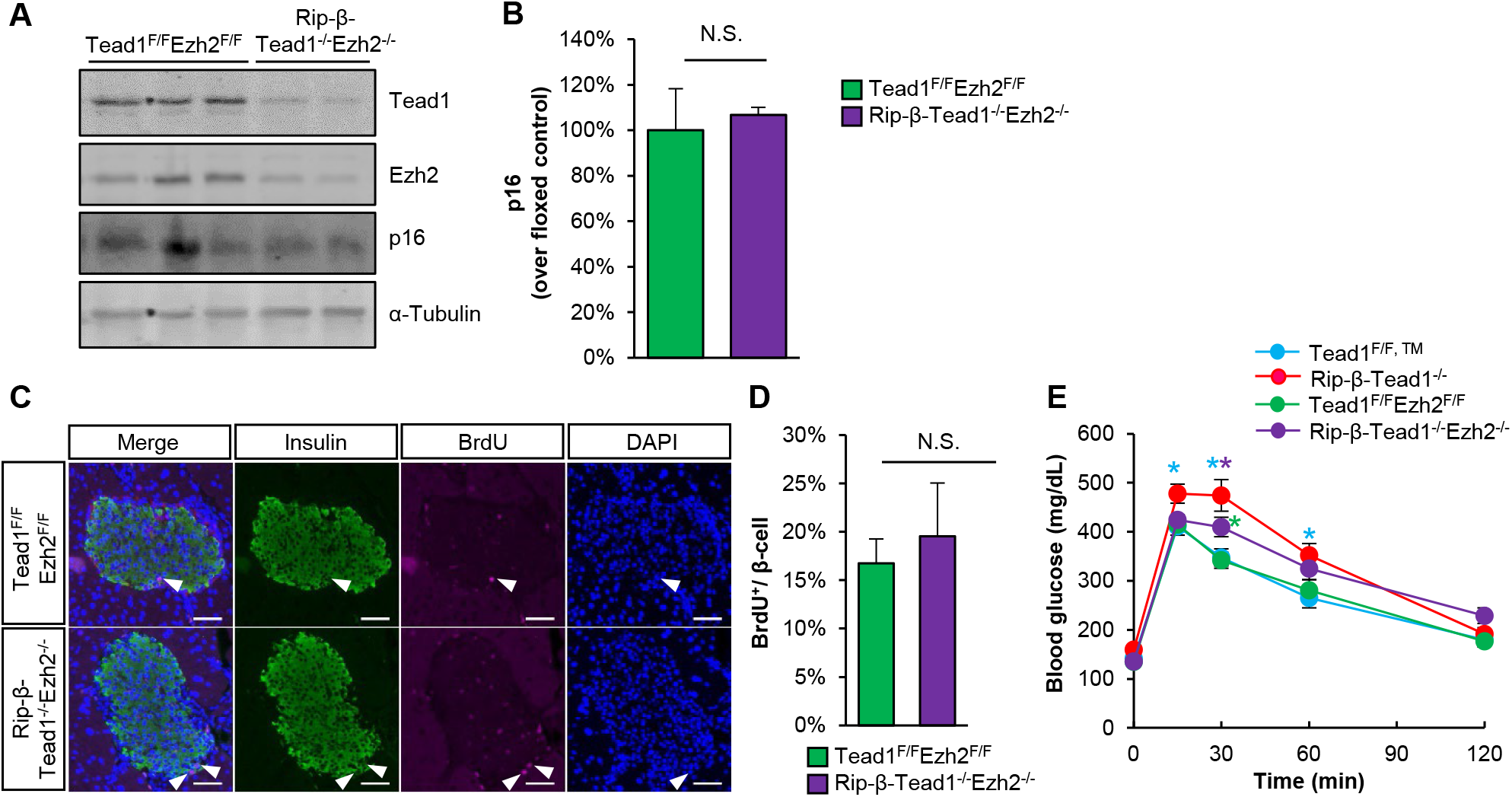
Tead1 regulates proliferative quiescence by its direct transcriptional regulation of p16^INK4a^ expression. (A) western blotting for Tead1, Ezh2 and p16 and (B) quantification of p16 protein in isolated islets from Tead1^F/F^Ezh2^F/F^ and Rip-β-Tead1^-/-^Ezh2^-/-^ mice. n=4-6. (C) Representative images of immunofluorescence staining for BrdU and insulin in Tead1^F/F^Ezh2^F/F^ and Rip-β-Tead1^-/-^Ezh2^-/-^ pancreas. BrdU was injected 2 hrs prior to sacrifice. (D) Quantification of β-cell proliferation in Tead1^F/F^Ezh2^F/F^ and Rip-β-Tead1^-/-^Ezh2^-/-^ mice at 10-11 weeks. n=3. (E) Blood glucose during GTT in overnight fasted 9-10 weeks old male mice. n=4-6.

## Discussion

Proliferative quiescence often accompanies the acquisition of mature function in many cell types, including β-cells ^5,47^. The molecular mechanisms that underpin this reciprocal regulation remain largely unexplored and this knowledge gap limits therapeutic approaches in regenerative medicine. Here, we identify Tead1 as mediating this reciprocal regulation of proliferative quiescence and maintenance of mature function in β-cells by direct transcriptional activation of a critical network of genes regulating these processes (schematically shown in **Figure 7**) in both mouse and human β-cells. We demonstrate that in adult β-cells Tead1 functions to limit proliferation and maintain mature identity and functional competence and its loss-of-function leads to β-cell failure and diabetes. Recent studies have highlighted limitations of individual insulin-promoter driven Cre-lines, such as potential secretory defects and hypothalamic expression in Rip-Cre, Gh minigene effects in Mip-Cre/ERT and possibility of low efficiency in Ins1-Cre and Mip-Cre/ERT lines ^30,48-50^. To overcome these limitations we have used a combination of multiple β-cell Cre-line derived genetic models to decipher the critical and non-redundant function that Tead1 exerts in adult β-cells.

**Figure 7:**
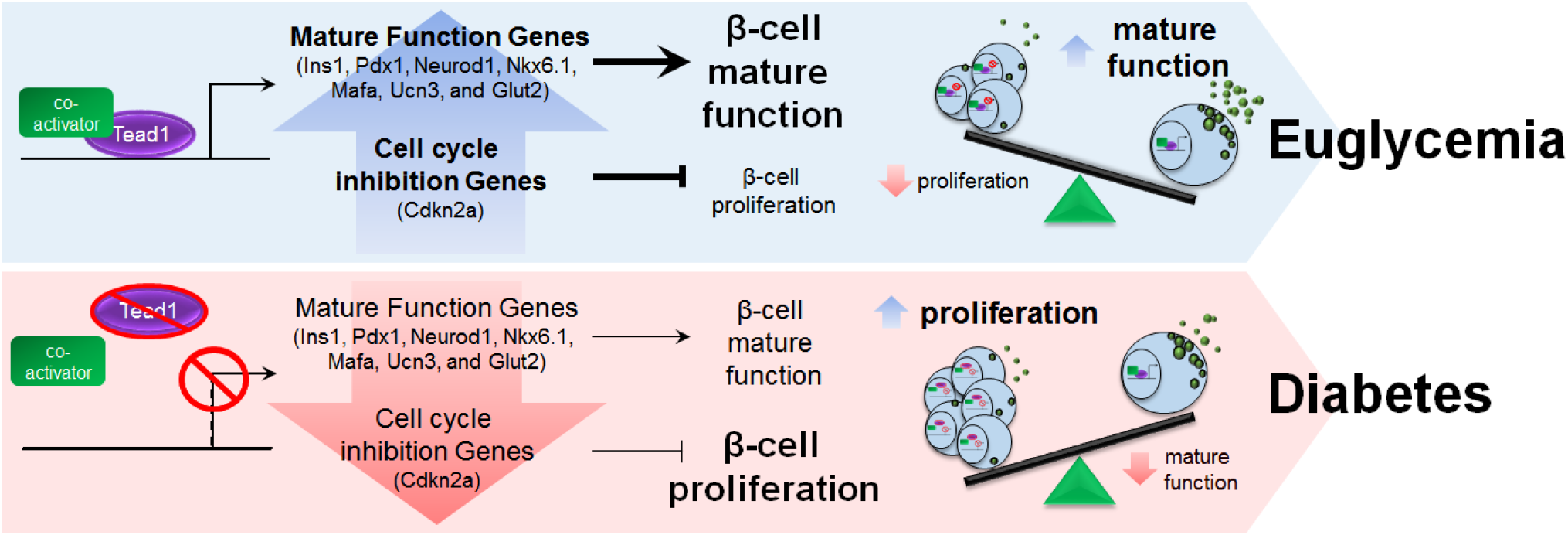
Schematic model of reciprocal regulation of β-cell proliferation and mature function by Tead1.

β-cells are highly proliferative before weaning, with proliferative rates as high as 4-6% ^51^. After weaning, they acquire GSIS with robust expression of gene networks regulating glucose sensing, transport, mitochondrial function, oxidative phosphorylation, and secretory machinery. However, as they gain functional competence, adult β-cells repress their proliferative program, lose their high basal proliferative rate having largely exited from cell cycle and self-renew at low rates that decline further with aging ^52-54^. While many pathways that regulate proliferation ^55-57^ and others that determine functional competence have been identified, mechanisms that evolved to coordinate these two programs remain poorly understood. In this study, we identify Tead1, as the critical mediator of this coordination of proliferative quiescence and functional competence in adult β-cells.

In the mature state, p16^INK4A^ (encoded by *Cdkn2a*) and its regulators in the polycomb repressive complex (PRC 1 and 2) are one of the primary enforcers of proliferative quiescence in adult β-cells ^44-46,58,59^. These exhibit a significant age-dependent regulation of proliferation ^41,44,45,59-62^. The unexpected finding that Tead1 is required for maintaining proliferative quiescence in mature β-cells led us to identify Tead1 as a novel transcriptional regulator of p16 and a loss of this regulation led to a reacquisition of a high proliferative rate. Deletion of Ezh2 to restore p16 expression in Tead1-deficient β-cells demonstrated that modulation p16 mediated the Tead1 enforced proliferative quiescence in mature β-cells. It is possible that Tead1 also interacts with other regulators of p16 including Ezh2, Bmi1 among others, especially in an age-related manner to restrict β-cell proliferation. In addition, concurrent work revealed that Tead1 also directly interacted with corepressors VGLL4 and MENIN to keep adult β-cells in proliferative quiescence^63,64^. How to modulate these Tead1 related regulatory mechanisms to enhance functional β-cell mass will be a subject for future studies.

Neonatal β-cells express many genes that are ‘disallowed’ in mature β-cells, including hexokinase1 (HK1), AldolaseB, Ldha, Acot7, MCT1 ^34-36,65^, many of which have been associated with immature function characterized by blunted GSIS. In rodent β-cells, high expression of a network of critical β-cell transcription factors, including Pdx1, Neurod1, Nkx6.1, a switch from MafB to high MafA expression along with and their target genes including Glut2, Glucokinase, Ucn3 and others, establishes functional competence and mature identity exemplified by robust GSIS ^32,66^. Repression of these genes is associated with a loss of mature function accompanied by a fetal/immature gene expression phenotype with expression of ‘disallowed ‘genes, with high MafB and loss of MafA expression. Interestingly, loss of Tead1 function leads to a decrease in expression of this gene network and loss of functional competence, but is not associated with re-expression of many of the disallowed genes, such as *Ldha*, nor a re-expression of MafB, hallmarks of de-differentiation to an immature state ^66^. This is in contrast to that seen in proliferating β-cells with activation of c-Myc ^5^ or Notch ^67^, suggesting that the loss of mature function with Tead1 deletion may be independent of the proliferation phenotype. Interestingly, the β-cell deletion of Eed, an interactor of Ezh2 in PRC2, was reported to have a severe β-cell phenotype ^59^ with βEedKO mice having significant diabetes along with a significant loss of mature identity and increased expression of immature markers, although no proliferative rates were reported. While there are similarities in the loss of mature identity in these studies, the salient distinction with loss of Tead1 function is that the markers of de-differentiation are not increased, supporting the assertion that Tead1 promotes maintenance of mature identity.

While the requirement of Tead1 for normal β-cell function is demonstrable in these studies, it would be interesting to speculate whether this is dysregulated in disease states. While factors that regulate upstream hippo kinase cascade can certainly play a role, other stimuli may also play a direct role in regulating Tead1 function, independent of the hippo kinase cascade. Recent studies have shown that Tead1 subcellular localization could be altered by p38 Map kinase activation and glucose itself ^68^ or its stability of co-factor binding can be regulated by palmitoylation ^69-71^. This raises the intriguing possibility that stress pathways and glucolipotoxicity could affect β-cell function via changes in Tead1 localization and activity. Could then glucotoxicity, lipotoxicity or other stress-induced induced β-cell dysfunction be modulated by regulation of Tead1 activity and provide novel therapeutic targets? Answers to these questions in future studies could uncover new druggable targets for therapy of diabetes.

This reciprocal regulation of mature function and proliferative quiescence enforced in β-cells, by Tead1, could underpin the hurdles in inducing β-cell proliferation without loss of function. Intriguingly, recent studies in human islets demonstrate that although YAP is not normally expressed in adult β-cells, its overexpression and activation of TEAD1 pathway could lead to proliferation without loss of function ^21,22^. This is also consistent with Tead1 regulation of PDX1 observed in this study. These observations suggest that Tead1 activity could be determined by the specificity of upstream coactivators and open the possibility that there are as yet undetermined pathways that are regulated by Tead1. While the regulators of Tead1 action in β-cells need further exploration, identification of these would allow therapeutic modulation of Tead1 activity to enhance β-cell replacement therapy.

In summary, Tead1 is a critical β-cell transcription factor required for coordinating adult β-cell proliferative quiescence, mature identity and functional competence to maintain glucose homeostasis. Modulation of this reciprocal regulation could help overcome hurdles in inducing β-cell proliferation without loss of function.

## Methods

### Animals

All animal experiments were approved by the Institutional Animal Care and Use Committee of Baylor College of Medicine and the University of Pittsburgh. Tead1 floxed ES cells were obtained from KOMP (Clone Name: EPDO207_2_H11), wherein the DNA binding domain coding Exon 3 is floxed. Targeted ES cells were injected into blastocysts to generate chimeric mice and those that demonstrated germline transmission were crossed with mice expressing FLP recombinase to remove the neomycin resistance cassette to generate Tead1 loxP alleles [floxed exon: ENSMUSE00000203030]. The homologous recombination was verified by long range PCR using primer set #1 (forward: GCAGTGCTCTCAGGAGTGCTGAGTGCGAC; reverse: AGAAGCCACAGTGCCCTGGAAGTGT) and primer set #2 (forward: GCCTTCTGAGTGCTGGCATTAAAGG; reverse: CACTGCAGCGCGCATAGCCTATGCTCCTTC) (**Figure S1**). Mice were genotyped using *Tead1*-flox-forward (GCCTTCTGAGTGCTGGCATTAAAGG) and reverse (AAGGCAGACTCCTTCATTGGAATGG) primers. Floxed Tead1 (Tead1^F/F^) mice were crossed with Rip-Cre or Ins1-Cre (Thorens) transgenic mice to generate β-cell-specific Tead1 deletion during embryonic development. For adult β-cell inducible deletion, Tead1 ^F/F^ mice were crossed with Mip-CreERT transgenic mouse (Ins1 promoter driven Cre-ER) to generate Tead1 ^F/F^ Mip-CreERT^+^ mice that when treated with Tamoxifen (TM) 100 mg/ kg every other day by gavage for 5 doses induced β-cell-specific Tead1 deletion (Mip-β-Tead1^-/-^). Ezh2 flox mice were obtained from Jackson labs (Stock#02616) and crossed into the Tead1^F/F^ background to obtain double floxed mice and crossed with Rip-Cre mice to obtain double knockout mice. Mice were maintained under standard 12-h light-dark cycles with ad lib access to food and water, unless specified otherwise.

### *In vivo* experiments

Glucose tolerance tests (GTTs) and acute insulin secretion tests were performed in 16 hour overnight fasted mice administrated 1.5 g/kg or 3 g/kg D-glucose i.p. respectively. Insulin tolerance test (ITT) was performed in 16 hour overnight fasted mice administrated 0.75 U/kg insulin i.p.

### Metabolic cage measurements

Metabolic cage measurements were performed as described previously ^72^. Oxygen consumption (VO_2_), energy expenditure, carbon dioxide production (VCO2), and the respiratory exchange ratio (RER) were measured with the comprehensive laboratory animal monitoring system (Columbus Instruments) in individual cages without bedding. Mice were acclimated to the metabolic cages for 4 days prior to the start of data collection. Data was collected for 72 h. All data was analyzed and averaged for the dark and light cycles separately.

### Islet isolation and *ex vivo* insulin secretion

Islets were isolated as previously described ^72,73^. Briefly, 1mg/ml collagenase P (Roche) dissolved in HBSS was injected into the common bile duct of anesthetized mice and the pancreas was removed, minced and incubated in collagenase solution at 37°C for 17 min. After three washes with 10% FBS, the islets were pelleted at 200 g and purified on Histopaque (Sigma) gradient. Islets from each mouse were individually purified by handpick for all experiments. The islets were incubated in 11.1 mM RPMI-1640 supplemented with 10% FBS and 1% penicillin/streptomycin, L-Glutamine overnight. GSIS studies were performed as described previously ^73^. Ten similar sized islets from individual mice were used, with at least three mice from each genotype, under each experimental condition. Incubation times for ex vivo insulin secretion studies were for 30 min for each condition and the results were normalized to the DNA content or to the total insulin content of the islets measured after acid ethanol extraction at the end of the experiment. All insulin assays were performed using mouse insulin ELISA kit (Mercodia). Insulin content was assayed after acid-ethanol extraction from whole pancreas and isolated islets and normalized to DNA and protein measured from the same extract. Similar protocol was used for the GSIS experiments in 832/13 cells.

### Gene and protein expression

Quantitative PCR (qPCR) from cDNA made from DNase-digested RNA was performed with gene-specific primers with SYBR green mix and normalized to the expression of housekeeping genes (*Tbp*, *TopI,* and *Actb*). The sequences of the primers used are available in Supplementary table 2.

Islets were isolated from each mouse individually as described above and 150–200 islets were snap frozen immediately. Western blotting was performed, as described before ^72^. Briefly, total protein was extracted from 150-200 islets mice using a protein lysis buffer with a cocktail of protease inhibitors and separated by a standard SDS-PAGE and transferred to a nitrocellulose membrane. Tead1 antibody (rabbit polyclonal, 1:10,000; Abcam), p16 antibody (mouse monoclonal, 1:1,000; Abcam), Pdx1 (rabbit polyclonal, 1:10,000; BCBC), Mafa antibody (rabbit polyclonal, 1:1,000; Bethyl lab Inc.), Ezh2 antibody (rabbit polyclonal, 1:1,000; Cell Signaling), α-tubulin (mouse monoclonal, 1:10,000; Cell Signaling), Hsp90 (mouse monoclonal, 1:1,000; Santa Cruz) and β-actin antibody (rabbit monoclonal-horseradish peroxidase conjugate, 1:10,000; Cell Signaling), as a housekeeping control, were visualized by enhanced chemiluminescence (Pierce).

### Immunostaining

Mouse pancreas were harvested, embedded in paraffin and sectioned to 5 µm thickness. Immunostaining was performed as previously described ^72^. Primary antibodies used were Glut2 (rabbit polyclonal, 1:200; Millipore), Insulin (guinea pig ployclonal, 1:500; Abcam), Glucagon (rabbit polyclonal, 1:400; Dako), Tead1 (rabbit polyclonal, 1:200; Abcam), Taz (rabbit polyclonal, 1:200; Santa Curz), Yap (rabbit poly clonal 1:200; Cell signaling), Pdx1 (guinea pig polyclonal, 1:500; Abcam), Mafa (rabbit polyclonal, 1:100; Bethyl lab), Ucn3 (rabbit polyclonal, 1:2,000; MGI), Ki67 (rabbit poly clonalm 1:50; Abcam), Brdu (rat polyclonal, 1:1,000; Abcam), Caspase-3 (rabbit polyclonal, 1:200; Cell signaling), and Cleaved Caspase-3 (rabbit monoclonal, 1:200; Cell signaling). To measure total pancreatic islet area, pancreas paraffin blocks from mice were cut 5 µm thickness sections spaced 150 µm apart between each slide. 5 pancreatic sections per mouse were stained and measured islets area using ImageJ software. For assessing number of Brdu positive and Ki67 expression cells in β-cell, 3,000 to 5,000 cells were counted from mouse pancreatic sections in 3 to 4 mice from each group.

### Cell culture and human islet

832/13 (Ins-1, courtesy of Dr. Christopher Newgard) rat β-cell line was cultured as described previously ^72^. Ins-2 murine β-cell line (Courtesy of Akio Koizumi) ^74^ was cultured in high glucose DMEM (4.5 g/L) containing 15% fetal bovine serum, 150 μM beta-2-mercaptoethanol, and Penicillin (100 units/ml)/Streptomycin (100 μg/ml). EndoC-β H2 (purchased from EndoCells and courtesy of Drs. Ravassard and Scharfmann) ^75^ were cultured according to the manufacturer’s instruction. EndoC-βH2 cells were cultured in DMEM (1 g/L Glucose) containing 2% BSA fraction V, 50 μoM 2-mercaptoethanol, 10 mM nicotinamide, 5.5 μg/ml transferrin, 6.7 ng/ml sodium selenite, Penicillin (100 units/ml)/Streptomycin (100 μg/ml). Tissue culture flasks and dishes were coated with DMEM (4.5 g/L Glucose) containing fibronectin (2 μg/ml), 1 % ECM, and Penicillin (100 units/ml)/Streptomycin (100 μg/ml). De-identified human islets were obtained from the Integrated Islet Distribution Program (IIDP). Donor characteristics are shown in Supplementary table 3. Received human islets were selected by hand picking and cultured in CMRL1066 (Mediatech, Corning) supplemented with 10% FBS, 1% P/S and 2 mM L-glutamine. Intact islets were used for Verteporfin (VP) treatment experiments. For adenovirus (AdV) infection, human islets were disassociated with 0.05% Trypsin as previously described ^76^ and subjected to infection at 20 MOI, the islets re-aggregated spontaneously and formed pseudoislets in 2 days. Protein samples were collected post 72hrs of AdV infection. All cell lines were initially authenticated by the donating investigator or vendor and were tested to be mycoplasma free.

### *In vitro* experiments

Lentiviral shRNA against Tead1 and scrambled shRNA controls were obtained from Thermo Scientific and used to generate stable cell lines with mouse insulinoma (832/13 and Ins-2) cells. The following shRNAs were used: shRNA-Tead1#1: 5’-CTCGCCAATGTGTGAATATA-3’, shRNA-Tead1#3: 5’-GACTGAACCTGGTTAATTTA-3’. The knockdown efficiency was confirmed by RT-qPCR and western blotting.

Native *Pdx1* and *Cdkn2a* promoter containing Tead1 binding element were amplified by PCR, cloned to pGL3 basic, mouse insulinoma cell line Ins-2 was used for the promoter activity assay.

Chromatin immunoprecipitation (ChIP) was performed using standard protocols in Ins-2 cells with Tead1 and Myc antibody or control IgG for pulldown. Immunoprecipitated DNA was then used as the template for qPCR with primers flanking conserved Tead1 binding element on promoters of target genes.

### Microarrays

Biotin-labeled aRNA was generated from total RNA from isolated islets from 3 mice of each genotype at 12 weeks of age and was tested for integrity on the Agilent Bioanalyzer and quantified using the NanoDrop® ND-1000 spectrophotometer. 12 μg of labeled aRNA samples were fragmented and re-checked for concentration and size. Hybridization cocktails containing Affymetrix spike-in controls, 12 μg of each fragmented labeled aRNA and the Arcturus Turbo Blocking Reagent heated to 99 °C for 5 minutes, then incubated at 45 °C for 5 minutes. The hybridization cocktails were loaded onto Affymetrix GeneChip Mouse 430 2.0 arrays. The arrays were hybridized for 16 hours at 45 °C with rotation at 60 rpm in the Affymetrix GeneChip® Hybridization Oven 640.The arrays were washed and stained with a streptavidin, R-phycoerythrin conjugate stain using the Affymetrix GeneChip® Fluidics Station 450. Signal amplification was done using biotinylated antistreptavidin.The stained arrays were scanned on the Affymetrix GeneChip® Scanner 3000. The images were analyzed and quality control metrics recorded using Affymetrix Command Console software version 3.0.0. The data was normalized by quantiles and then significant genes analyzed using multiple sample corrected t-testing and 1.5 fold change and by gene ontology terms and MSigDb searches.

### Rna-Seq

Total RNA samples with RNA integrity number (RIN) ≥8 were used for transcriptome sequencing. Total RNA (10ng) from a pooled sample from 1 year old mice (n=3) each for two independent samples for each group was used for amplified double-stranded cDNA that was sheared to 200-300bp and ligated to Illumina paired-end adaptors using the Illumina TruSeq DNA library preparation kit according to the manufacturer’s instructions (Illumina, San Diego, CA). PCR amplification was performed to obtain the final cDNA library using Illumina kits. Bioanalyzer 2100 (Agilent Technologies, Santa Clara, CA) analysis was used to verify fragment size after amplification, library size and concentration before clustering. A total of 10pM of the library was then used for paired-end sequencing on the HiSeq 2500 at the Sequencing core at Brigham Young University. All further analysis were performed using the CLC genomics workbench ver 12. Raw RNA sequencing reads were mapped to the mouse reference genome build 10 (UCSCmm10/GRCm38). Mapped reads were counted using the feature counts and differential expression between the samples analyzed using multiple hypothesis testing. Pathway analysis: Gene Set Enrichment (GSEA, version 2.2.3, http://software.broadinstitute.org/gsea/) was performed on normalized count per million (CPM) or on ranked gene lists. Ranked lists were created based on the expression levels of DEGs in Rip-Tead1 KO vs. Flox controls (q<0.05). Significance was assessed by analyzing signal-to-noise ratio and gene permutations based on 1,000 permutations. Molecular signature database (MSigDB) 3.0 curated gene sets for hallmark and canonical pathways were used for the analysis. Significant gene sets with enrichment score and a q value cutoff of 0.05 are presented.

### ATAC-Seq

Flash-frozen tissue was sent to Active Motif to perform the ATAC-seq assay. The tissue was manually disassociated, isolated nuclei were quantified using a hemocytometer, and 100,000 nuclei were tagmented as previously described (Buenrostro et al. 2013), with some modifications based on (Corces et al. 2017) using the enzyme and buffer provided in the Nextera Library Prep Kit (Illumina). Tagmented DNA was then purified using the MinElute PCR purification kit (Qiagen), amplified with 10 cycles of PCR, and purified using Agencourt AMPure SPRI beads (Beckman Coulter). Resulting material was quantified using the KAPA Library Quantification Kit for Illumina platforms (KAPA Biosystems), and sequenced with PE42 sequencing on the NextSeq 500 sequencer (Illumina). Analysis of ATAC-seq data was very similar to the analysis of ChIP-Seq data. Reads were aligned using the BWA algorithm (mem mode; default settings). Duplicate reads were removed, only reads mapping as matched pairs and only uniquely mapped reads (mapping quality >= 1) were used for further analysis. Alignments were extended in silico at their 3’-ends to a length of 200 bp and assigned to 32-nt bins along the genome. The resulting histograms (genomic “signal maps”) were stored in bigWig files. Peaks were identified using the MACS 2.1.0 algorithm at a cutoff of p-value 1e-7, without control file, and with the –nomodel option. Peaks that were on the ENCODE blacklist of known false ChIP-Seq peaks were removed.

### ChIP-seq

Frozen tissue was sent to Active Motif Services (Carlsbad, CA) to be processed for ChIP-Seq. In brief, tissue was submersed in PBS + 1% formaldehyde, cut into small pieces and incubated at room temperature for 15 minutes. Fixation was stopped by the addition of 0.125 M glycine (final). The tissue pieces were then treated with a TissueTearer and finally spun down and washed 2x in PBS. Chromatin was isolated by the addition of lysis buffer, followed by disruption with a Dounce homogenizer. Lysates were sonicated and the DNA sheared to an average length of 300-500 bp. Genomic DNA (Input) was prepared by treating aliquots of chromatin with RNase, proteinase K and heat for de-crosslinking, followed by ethanol precipitation. Pellets were resuspended and the resulting DNA was quantified on a NanoDrop spectrophotometer. Extrapolation to the original chromatin volume allowed quantitation of the total chromatin yield. An aliquot of chromatin (10 ug for islets and 30 ug for hearts) was precleared with protein G agarose beads (Invitrogen). Genomic DNA regions of interest were isolated using 20 ul of antibody against Tead1. Complexes were washed, eluted from the beads with SDS buffer, and subjected to RNase and proteinase K treatment. Crosslinks were reversed by incubation overnight at 65 C, and ChIP DNA was purified by phenol-chloroform extraction and ethanol precipitation. Quantitative PCR (QPCR) reactions were carried out in triplicate on specific genomic regions using SYBR Green Supermix (Bio-Rad). The resulting signals were normalized for primer efficiency by carrying out QPCR for each primer pair using Input DNA. Illumina sequencing libraries were prepared from the ChIP and Input DNAs by the standard consecutive enzymatic steps of end-polishing, dA-addition, and adaptor ligation. Steps were performed on an automated system (Apollo 342, Wafergen Biosystems/Takara). After a final PCR amplification step, the resulting DNA libraries were quantified and sequenced on Illumina’s NextSeq 500 (75 nt reads, single end). Reads were aligned to the mouse genome (mm10) using the BWA algorithm (default settings). Duplicate reads were removed and only uniquely mapped reads (mapping quality >= 25) were used for further analysis. Alignments were extended in silico at their 3’-ends to a length of 200 bp, which is the average genomic fragment length in the size-selected library, and assigned to 32-nt bins along the genome. The resulting histograms (genomic “signal maps”) were stored in bigWig files. Peak locations were determined using the MACS algorithm (v2.1.0) with a cutoff of p-value = 1e-7. Peaks that were on the ENCODE blacklist of known false ChIP-Seq peaks were removed.

### Statistical methods

All statistical testing was performed either by two-tailed Student’s t-test, assuming unequal variance for two groups or ANOVA for multiple samples with p ≤ 0.05 considered significant between the groups.

Expression data have been deposited in GEO: 12 weeks (GSE139228) and 1 year (GSE139152).

Chip-seq and ATAC-seq data have been deposited in GEO under GEO accession GSE157513

## Acknowledgements

This work was supported by grants from the VA-ORD-BLR&D, I01BX002678 (V.Y.) and National Institutes of Health R01 DK097160 and DK130499 (V.Y.), National Institutes of Health K08-HL091176 and R01-HL147946 (M.M.), American Heart Association Career Development Award 19CDA34770034 (R.L.), National Institutes of Health CA125123 (C.J.C.) by the Human Tissue Acquisition and Pathology (HTAP) core (NIH; NCI P30-CA125123), the Integrated Microscopy Core (NIH-HD007495, DK56338, and CA125123, the Dan L. Duncan Cancer Center, and the John S. Dunn Gulf Coast Consortium for Chemical Genomics), the Mouse Metabolic Core (P30-DK079638), by the Genomic and RNA Profiling Core (P30-DK076938) at Baylor College of Medicine.

## Author Contributions

J.L. designed and conducted the experiments, performed data analyses, interpreted data and helped write the manuscript; R.L., B.S.K. helped with specific experiments and data analysis. P.K.S., O.S., M.O.H., F.L., R.J., P.Y., V.N., E.M.P-G., H.S., R.B., Y.Z., C.C., C.J.C., K.M., and M.M. provided critical scientific input, data analysis, and logistical support. R.L., M.O.H, K.M. and M.M. provided critical input into editing the manuscript. V.K.Y supervised the project, designed experiments, interpreted data and wrote the manuscript.

## Supplementary Figure Legends

**Supplemental Fig S1:**
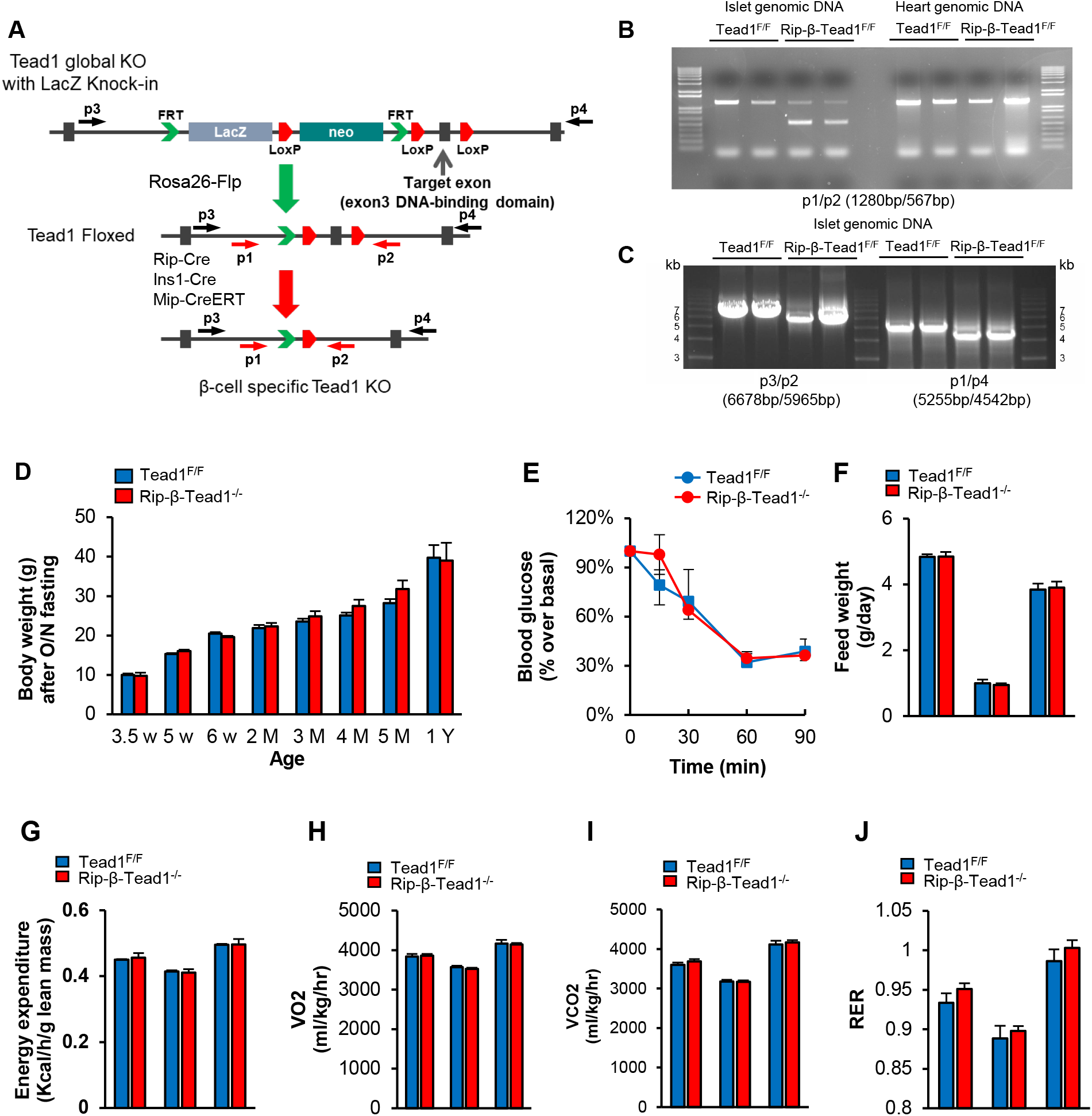
Generation of Tead1 floxed mouse strain and β-cell specific Tead1 knock out mice. (A) Generation of Tead1 conditional knockout allele, and β-cell specific knockout mouse models. (B) Genomic PCR showing the recombined fragment of the Tead1 allele in islet and heart from Tead1^F/F^ and Rip-β-Tead1^-/-^mice. (C) Tead1 flox allele and its deletion in Rip-β-Tead1^-/-^, verified by long-range PCR (right) in Tead1^F/F^ and Rip-β-Tead1^-/-^islet. (D) Body weight of Tead1^F/F^ and Rip-β-Tead1^-/-^mice. n=4-6. (E) Plasma glucose represented as a percent drop from baseline during insulin tolerance test in 12 weeks old male mice. n=5. All values are mean ± SEM. (F) Food intake measured over a 3-day period. n=4. Energy expenditure and VO2 (G and H), VCO2 (I), and RER (J) were assessed as described in Methods in supplementary material. n=4. All values are mean ±SEM.

**Supplemental Fig S2:**
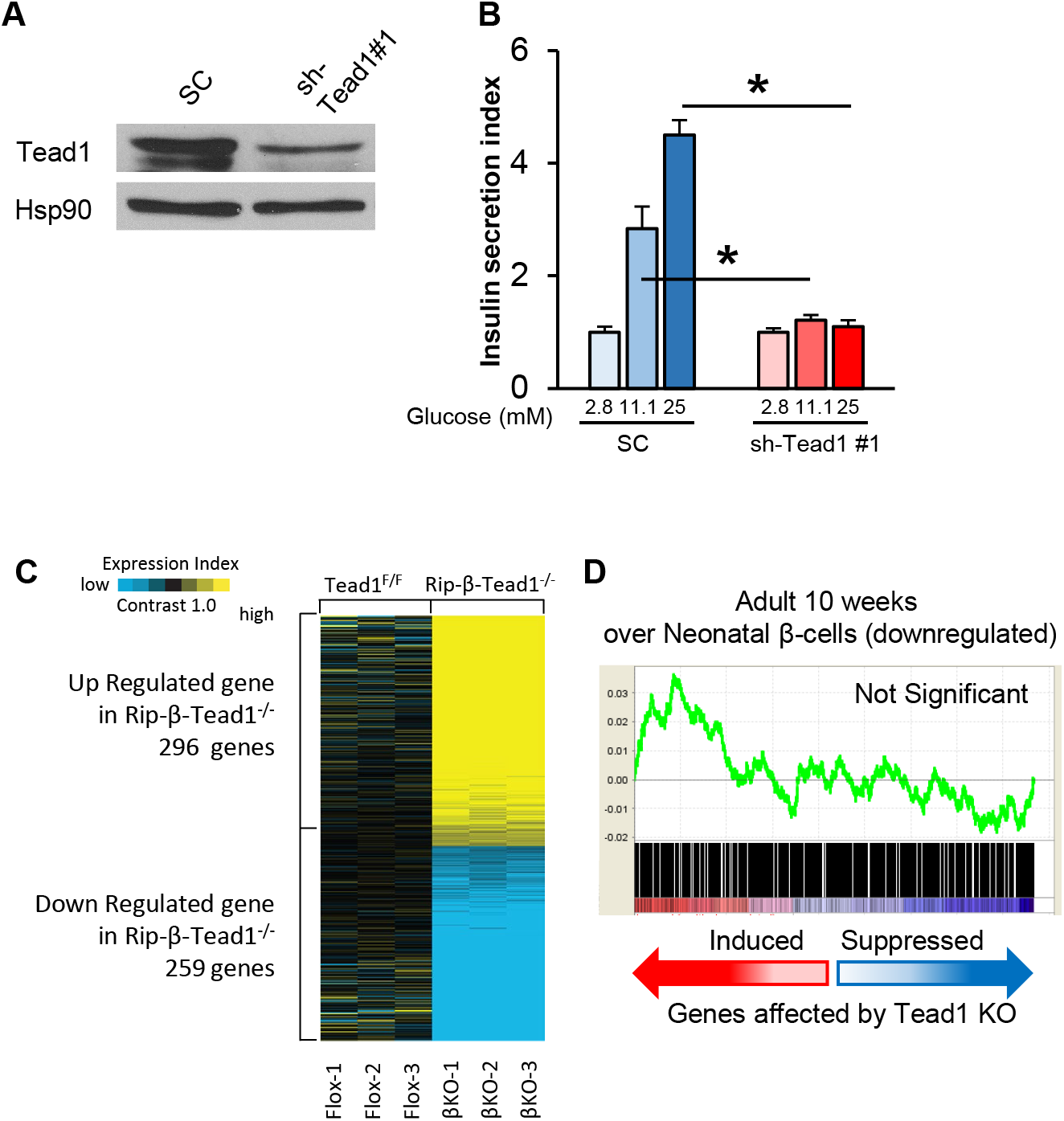
Cell-autonomous regulation of mature β-cell function by Tead1. (A,B) Tead1-depleted β-cells have impaired GSIS in vitro. (A) Western blotting for Tead1 in shRNA Tead1 knockdown and scrambled Ins-1 (832/13) cells. (B) GSIS in response to increasing glucose concentration in Tead1-knockdown and scrambled control 832/13 cells. n= 4. All values are mean ± SEM, * - p≤ 0.05. (C) Transcriptome analysis heat map showing up and downregulated genes in isolated islets from 12 week old Rip-β-Tead1^-/-^ compared to Tead1^F/F^ mice. (D) GSEA analysis of the differentially expressed upregulated genes in isolated islets of Rip-β-Tead1^-/-^ compared to Tead1^F/F^ as compared to the published gene set (GSE47174) that was upregulated in adult β-cells when compared to neonatal β-cells. Details in the accompanying text. FDR Q value did not show any significance.

**Supplemental Fig S3:**
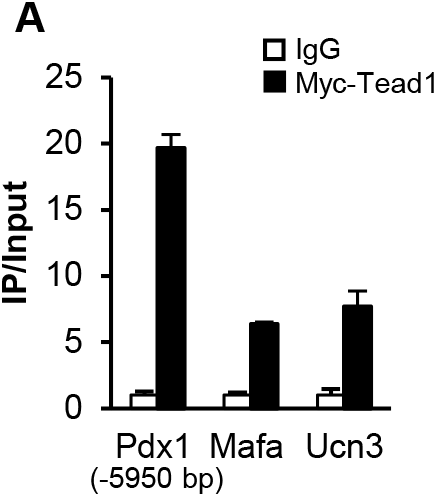
Tead1 regulates mature β-cell function-related genes. (A) ChIP of Myc-Tead1 over-expressing Ins-2 cells with Myc or control IgG antibody and qPCR with primers flanking putative Tead1 response elements in promoters of displayed genes. The y-axis represents the ratio of pulldown DNA to input DNA. n=3.

**Supplemental Fig S4:**
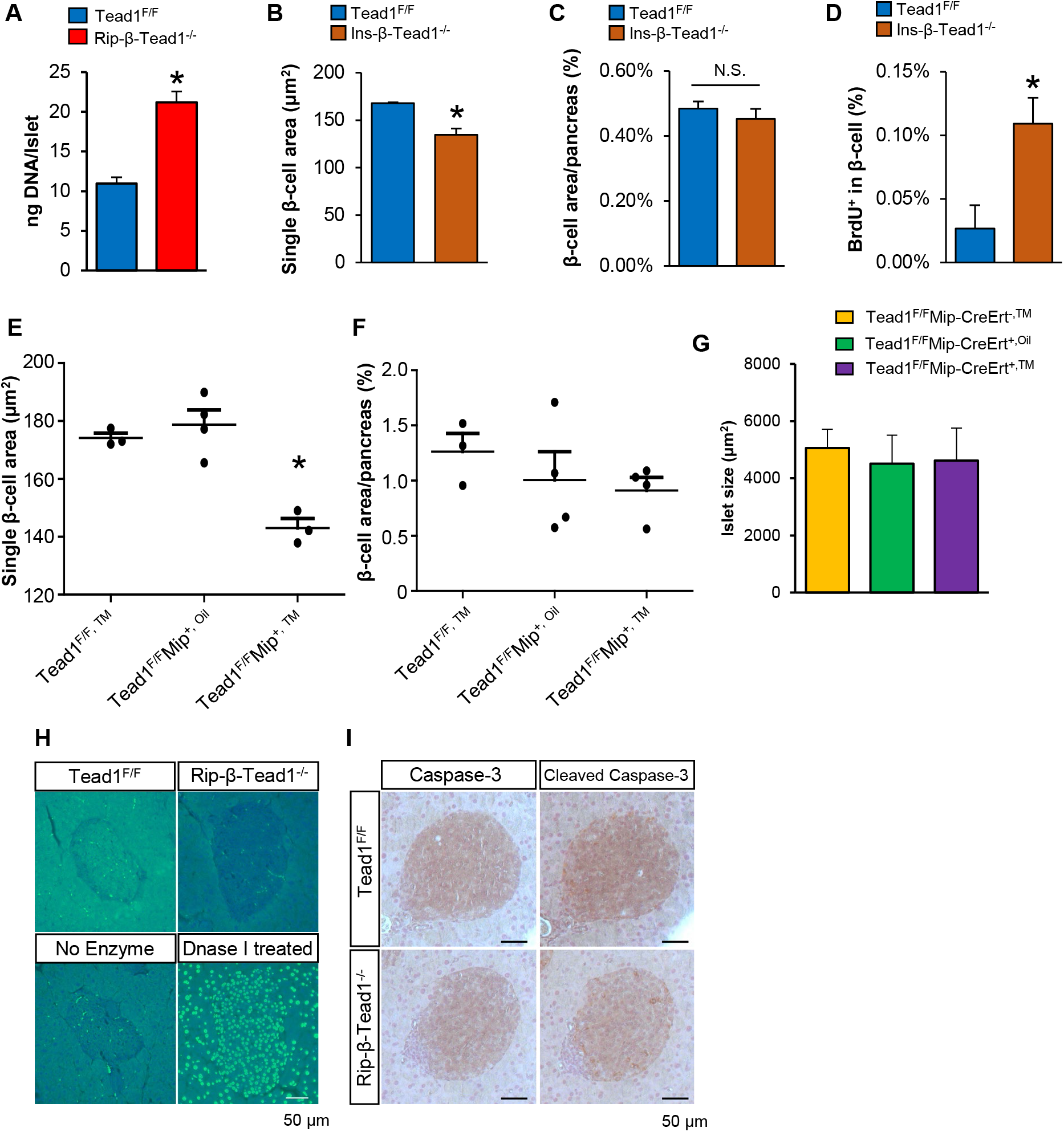
Tead1 regulates β-cell proliferative quiescence. (A) Genomic DNA content in similar sized islets from Rip-β-Tead1^-/-^ and floxed control mice. n=5. (B,C, and D) Quantification of (B) β-cell size, (C) β-cell area and (D) BrdU positive β-cells in 5-6 month old Ins-β-Tead1^-/-^ and floxed control mice n=3-4. (E, F, and G) Quantification of (E) β-cell size, (F) β-cell area and (G) average islet size in Mip-β-Tead1^-/-^ and control mice n=3-4. (H and I) Apoptosis is not altered in Tead1-deficient β-cells. Representative image of (H) TUNEL staining and (I) immunohistochemistry for Caspase-3 and cleaved Caspase-3 in Rip-β-Tead1^-/-^ and control islet. DNase I treatment was used as a positive control for TUNEL staining. Scale bar – 50μm.

